# Chromatin co-accessibility is highly structured, spans entire chromosomes, and mediates long range regulatory genetic effects

**DOI:** 10.1101/604371

**Authors:** William W. Young Greenwald, Agnieszka D’Antonio-Chronowska, Paola Benaglio, Hiroko Matsui, Erin N. Smith, Matteo D’Antonio, Kelly A. Frazer

## Abstract

Chromatin accessibility identifies active regions of the genome, often at transcription factor (TF) binding sites, enhancers, and promoters, and contains regulatory genetic variation. Functionally related accessible sites have been reported to be co-accessible; however, the prevalence and range of co-accessibility is unknown. We perform ATAC-seq in induced pluripotent stem cells from 134 individuals and integrate it with RNA-seq, WGS, and ChIP-seq, providing the first long-range chromosome-length analysis of co-accessibility. We show that co-accessibility is highly connected, with sites having a median of 24 co-accessible partners up to 250Mb away. We also show that co-accessibility can *de novo* identify known and novel co-expressed genes, and co-regulatory TFs and chromatin states. We perform a *cis* and *trans*-caQTL, a *trans-*eQTL, and examine allelic effects of co-accessibility, identifying tens of thousands of *trans*-caQTLs, and showing that *trans* genetic effects can be propagated through co-accessibility to gene expression for cell-type and disease relevant genes.

## Introduction

Regulatory genetic variation that affects gene expression and human disease is often found within accessible chromatin sites^1-6^. These accessible sites, measured by either DNase-seq^7,8^ or ATAC-seq^9,10^, identify functional regions of the genome including active promoters and enhancers^8,11,12^, as well as the transcription factor (**TF**) binding sites within them^8,13-15^. However, it is difficult to determine the function of accessible sites as they can be distal from their targets^1^. In order to identify the functionality of accessible sites, previous studies have examined co-accessibility: the coordination of specific chromatin accessibility sites. These studies have examined co-accessibility at fine-scale (ie specific accessible sites) for local *cis* interactions within 10kb (ie co-binding TFs, promoter regulation, and local enhancer regulation)^5,6,16^, and long-range *cis* interactions between 10kb and 1.5Mb (i.e. chromatin looping and distal enhancer regulation)^5,16^. They have mainly applied supervised approaches to show that co-accessibility occurs between regulatory regions and their targets, as well as co-binding TFs, and that the majority of *cis* acting, genetically associated co-accessibility occurs at sites <20kb apart^16^. However, due to computational and statistical power, these studies limited their examination of co-accessibility either to fine-scale resolution and local structure, or higher order properties across long-ranges at low resolution^5^. It is thus unclear if co-accessibility extends to *cis* (ie physical co-regulation such as a TF co-binding or looping) and *trans* (ie sequential co-regulation such as a gene network) relationships across long distances (10s-100s of megabases), how many sites across a chromosome are co-accessible with one another (ie how highly connected is co-accessibility), and whether accessible sites can mediate genetic effects on highly distal sites via co-accessibility. A more comprehensive understanding of the co-accessible chromatin landscape and its genetic associations could provide novel insights into the effects of regulatory genetic variation across short and long distances.

As gene regulation involves many distal regulatory components, it is expected that genetic variation could exert *trans* long range regulatory effects. The omnigenic model^17^ of gene regulation has recently estimated that 70% of the heritability of gene expression is due to *trans* effects^18^. However, these genetic effects are thought to have effect sizes orders of magnitude smaller than *cis* effects^18^, and as they are distal from their targets, they are extremely difficult to identify (creating a power problem due to multiple-testing burden) and delineate from confounded *cis* effects. One possible solution to this statistical power problem could be to leverage chromatin co-accessibility to reduce search space, as gene expression and accessibility of the gene’s promoter are known to be correlated^19^. To overcome confounded *cis* effects, it could be possible to use mediator analyses in which one specifically tests for an intermediate effector rather than two independent associations. Additionally, studies examining chromatin accessibility quantitative trait loci (caQTLs) have found moderate overlap between *cis-*caQTLs and *cis-*eQTLs (∼30-40%)^3,5^. Thus, it is possible that co-accessibility could be used to tie regulatory elements to their distal co-regulators or gene targets, and then subsequently identify *trans* genetic effects. As this strategy would greatly reduce the number of variant-target pairs tested for *trans* effects (thus reducing multiple testing burden), it may be possible to observe hundreds or thousands of more *trans* effects than previous studies. Identifying co-accessible chromatin regions across entire chromosomes, and the genetic variation associated with these accessible regions, could therefore better elucidate the extent to which genetic variants exert long range *trans* effects, and how these effects may be mediated via co-accessibility.

Here, we perform ATAC-seq in 152 induced pluripotent stem cells (iPSCs) from 134 individuals from iPSCORE^20-23^, and integrate this data with available WGS and RNA-seq for the same individuals. We call over 1 million accessible chromatin sites and utilize population-level information to identify co-accessible sites by testing for correlation in accessibility between all sites chromosome-wide. We show co-accessibility is highly connected, with sites being co-accessible with an average of 24 other sites, and can span long distances (up to hundreds of megabases). We then use these significant relationships to create co-accessibility networks, and show that neighbors in these networks are enriched for TF co-binding partners, functionally related TFs, spatially colocalized loci (ie loci in a chromatin loop), and co-expressed genes up to 100Mb apart, and can also be used to infer novel TF functionality. Next, we examine the genetic architecture of co-accessibility by measuring allele specific effects (ASE) and performing one of the largest caQTLs studies to date. We show that genetic effects spread through co-accessibility, with highly connected sites being more likely to have a *cis-*caQTL or exhibit ASE; additionally, strong ASE explains 52% of co-accessible weaker ASE. Finally, we leverage these networks to identify more than 92,000 *trans-*caQTLs greater than 1.5Mb from their target, 9 of which are also *trans-*eQTLs for cell type and disease relevant genes. Overall, our data reveals that chromatin co-accessibility is highly connected, spans the length of entire chromosomes, can *de novo* identify co-regulatory TFs, is a mechanism underlying *trans* genetic effects, and can give insight into *trans-*eQTL mechanisms.

## Results

### Samples, ATAC-seq data generation, and ATAC peak characterization

To measure chromatin co-accessibility, accessible sites were identified from ATAC-seq performed on 152 iPSC lines. These lines were generated from 134 individuals (Supplementary Table 1) from iPSCORE and have previously been shown to be pluripotent and to have high genomic integrity^20^ (Figure 1A). We obtained a total of 5.5 billion reads, and after QC, filtering, and merging individual samples (see methods; Supplemental Table 1), inspected the quality of this data by examining its overlap and consistency with higher order chromatin structure at low-resolution, chromatin states, and H3K27ac peaks. To examine higher order structure (Figure 1B), we compared the correlation between ATAC-seq signal in 500kb bins across chromosome 18 to the correlation in Hi-C (from iPSCORE iPSCs^24^), and observed a similar pattern between the two as previously reported^5^. We next used MACS2 to call ATAC-seq peaks (obtaining a total of 1.01 million peaks), and examined the overlap of chromatin states from the iPSC ROADMAP^12^ with the peaks. We found peaks to be enriched for active TSS, transcribed regions, enhancers, polycomb-repressed, bivalent TSS, and bivalent enhancers, and depleted for repressed chromatin (heterochromatin and quiescent chromatin, Figure 1C). These findings are consistent with properties of accessible chromatin and known specialized use of bivalent and polycomb chromatin in maintaining iPSC pulirpotency^25-28^. Next, we examined the distribution of ATAC-seq reads at H3K27ac peaks from iPSCORE iPSCs and observed an enrichment at the centers of these H3K27ac peaks (Figure 1D). Together, these results show that this ATAC-seq data follows known characteristics of accessible chromatin and cell type specific characteristics of iPSCs.

**Table 1.** *trans*-eQTL results:

| Variant | Summit | Gene ID | Gene Name | Beta | P | Q | Variant Summit Distance |
| --- | --- | --- | --- | --- | --- | --- | --- |
| chr11:85425376 | chr11:560507 | ENSG00000099849.10 | RASSF7 | -3.517668244 | 8.09E-16 | 8.29E-12 | 84864869 |
| chr1:18566851 | chr1:243418251 | ENSG00000054282.11 | SDCCAG8 | 0.219172129 | 2.14E-07 | 0.001098407 | 224851400 |
| chr16:2553486 | chr16:29754764 | ENSG00000260719.1 | AC009133.17 | -0.74599912 | 4.05E-06 | 0.013810673 | 27201278 |
| chr9:139667161 | chr9:140683708 | ENSG00000181090.13 | EHMT1 | 1.774032818 | 2.07E-05 | 0.049666034 | 1016547 |
| chr19:9999210 | chr19:4685575 | ENSG00000142002.12 | DPP9 | 0.476342794 | 3.43E-05 | 0.049666034 | 5313635 |
| chr19:13023591 | chr19:2458281 | ENSG00000176619.6 | LMNB2 | 0.51023325 | 3.86E-05 | 0.049666034 | 10565310 |
| chr9:80914060 | chr9:116263136 | ENSG00000138835.18 | RGS3 | -0.637221951 | 4.09E-05 | 0.049666034 | 35349076 |
| chr11:66141427 | chr11:72370422 | ENSG00000186642.11 | PDE2A | -0.974316619 | 4.17E-05 | 0.049666034 | 6228995 |
| chr17:65421333 | chr17:38258696 | ENSG00000126368.5 | NR1D1 | 0.909156673 | 4.36E-05 | 0.049666034 | 27162637 |

**Figure 1:**
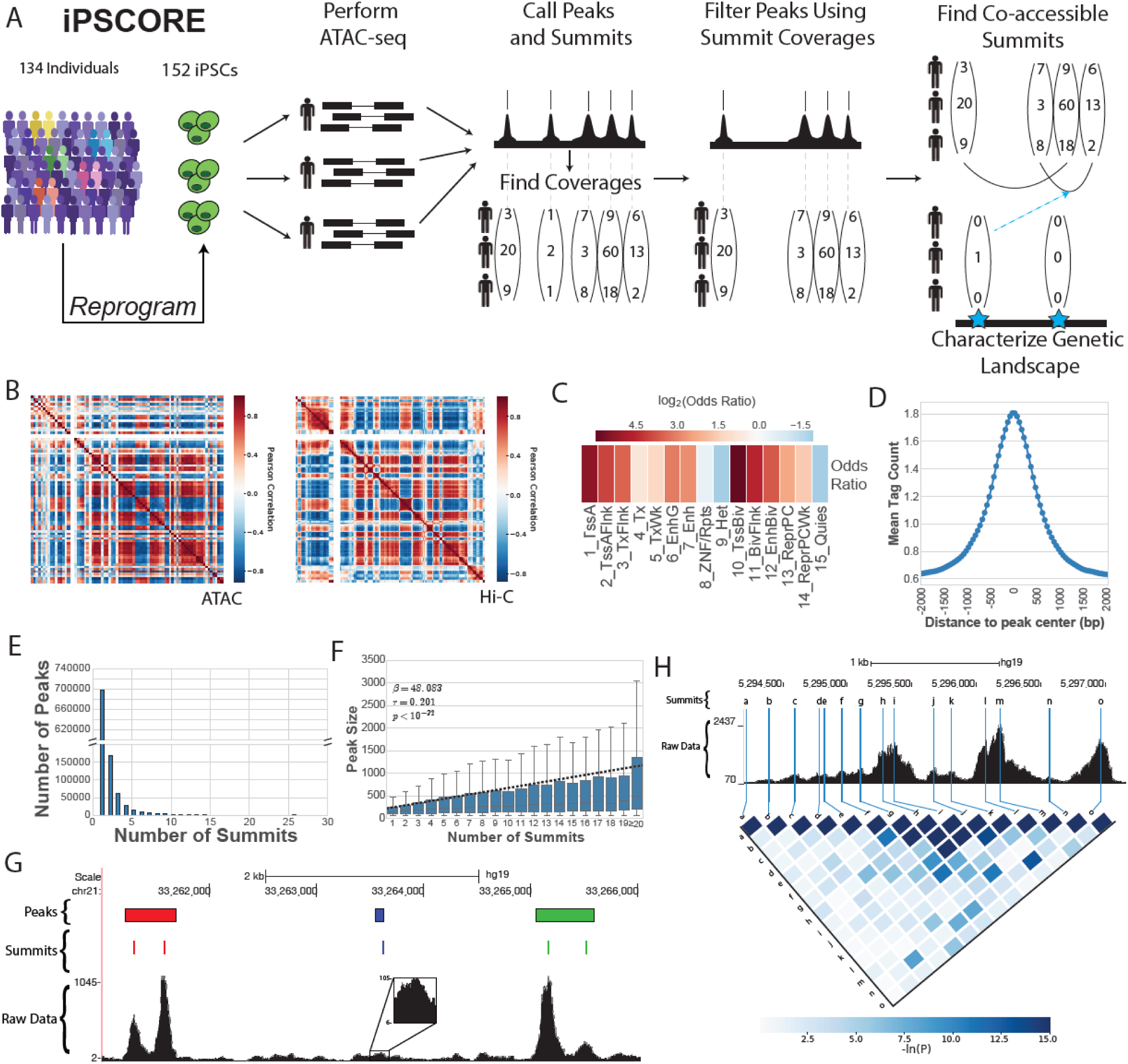
(A) Overview of experimental design. iPSCs from 134 individuals from the iPSCORE cohort were selected for ATAC-seq sequencing (152 samples total). After QC and filtering, all individual’s data was utilized to call peaks, and the coverages of each peak and summit were calculated. Peaks were subsequently filtered based on summit coverage, and co-accessible summits were found from summit data. Finally, genetic associations with co-accessibility were examined. (B) Correlation of total reads in 500kb bins for ATAC (left), compared to Hi-C Pearson matrix (right), across chromosome 18. Broadly similar patterns can be observed. (C) Heatmap of the log_2_(Odds Ratio) of chromatin state enrichments within ATAC-seq peaks relative to the genome, measured in number of base pairs in each state. Histogram of the average tag count of ATAC-seq reads at H3K27ac peaks across all samples. (E) Histogram showing the number of peaks that have a given number of summits. (F) Boxplot (middle two quantiles and median shown as box and line within box; outliers not shown) and regression line on raw data (dashed line) for the number of summits vs the length of the peak in base pairs. (G) Genome browser picture of the combined ATAC-seq data across all individuals (raw data), summit Calls (Summits), and peak calls (Peaks). Three peaks were called in this window, shown in red, blue, and green; summits are colored by the peak to which they belong. Both the red and green peak calls contain two seemingly distinct peaks, which were identified by their summit calls. The blue peak, while lower, is still peak-shaped and has high coverage (105 reads). (H) Genome browser picture for the combined ATAC-seq data across all individuals (raw data) at a single peak call with 15 summits (labeled a-o). Lines for summits are extended through the raw data, and connect to their label on the heatmap. Heatmap shows the negative natural log of the p-value of correlation between these summits. Correlation quickly decays as a function of distance, and notably, most summits are not strongly correlated.

### Peaks contain multiple non-co-accessible summits

As part of ATAC-seq data processing, reads are used to call peaks and their sub-peak structure (**summits**). As summits represent individual TF binding sites within peaks (https://github.com/taoliu/MACS), we examined the number of summits within peaks and found that while the majority of peaks contained a single summit, 31% (315,901) contained multiple summits, with some containing up to 26 (Figure 1E). Additionally, we found a strong relationship between the length of peak and the number of summits identified (Pearson Correlation p < 10^-32^; Figure 1F). These patterns are consistent with MACS2’s documentation (https://github.com/taoliu/MACS) stating that nearby individual binding sites are called as summits and binned together as a single peak call (Figure 1G). We tested whether the summits acted independently by examining the correlation between summit heights in the same peak across individuals, and found that 97.5% of summits were not significantly correlated with other summits within the peak (see methods). Further, the significance of correlation between summits within the same peak decayed with distance (Figure 1H). Together, these data indicate that many peak calls contained multiple independent accessible sites; we therefore utilized the 1.21 million summits ATAC summits from peaks that passed QC for downstream analyses of accessible chromatin sites.

### Co-accessibility is predominantly distal and highly connected, spanning entire chromosomes

We set out to characterize the local and long-range co-accessibility landscape at fine-scale resolution (ie site-by-site co-accessibility) for each of the 22 autosomes chromosome-wide. We tested the quantile normalized trimmed mean of M values (**TMMs**^**29**^) of coverage for each site with every other site pairwise on each chromosome using a Linear Mixed Model to account for covariates and kinship. For each pair, we obtained a regression coefficient (**β**) and a p-value. We performed FDR correction of the p-values by chromosome, obtaining between 45 thousand and 3 million significant co-accessible relationships per chromosome (FDR q < 0.05). We observed similar numbers of co-accessible pairs normalized to chromosome length, except for chromosome 19 (Supplemental Figure 1A) which was ∼4x higher. This increase may have been driven by a large cluster of Znf genes known to be highly coordinated^30^. We first examined the distribution of the number of sites each site is co-accessible with (**connectivity**) across all sites for each chromosome, and found the level of connectivity to vary (Figure 2A), ranging from sites with no co-accessible partners to those with ∼3,000, and a mean of 24.46 partners. Surprisingly, we found that 96.6% of all ATAC sites were co-accessible with at least one other site, suggesting that the vast majority of regulatory sites interact with at least one other regulatory site. We next measured the distances between co-accessible sites (Figure 2B). As expected, the most commonly observed distances (i.e. modes of the data) were within the ranges previously studied for local, likely *cis*, co-accessibility (<1.5 Mb Figure 2B). However, the vast majority of co-accessible sites were further than 1.5Mb apart, with some pairs extending up to 250Mb distal from one another and a mean distance of 48.94Mb. Additionally, we found the strength of association to be consistent across distances (Supplemental Figure 1B). Finally, to better understand these data, we visualized the specific site-by-site correlations across chromosome 18 (Figure 2C), and found that, as Figures 2A and 2B suggested, sites on opposing ends of the chromosome were co-accessible. Further, we zoomed in to eight different resolutions, and at each resolution, we found co-accessible sites spanning almost the entire window (Figure 2C). Together, these data indicate that fine-scale co-accessibility extends beyond the local *cis* structure of 1.5Mb that has been previously examined, to sites that are highly distal from one another and likely *trans* in nature (up to hundreds of Mb). Overall, these data reveal that co-accessibility is highly distal and inter-connected.

**Figure 2:**
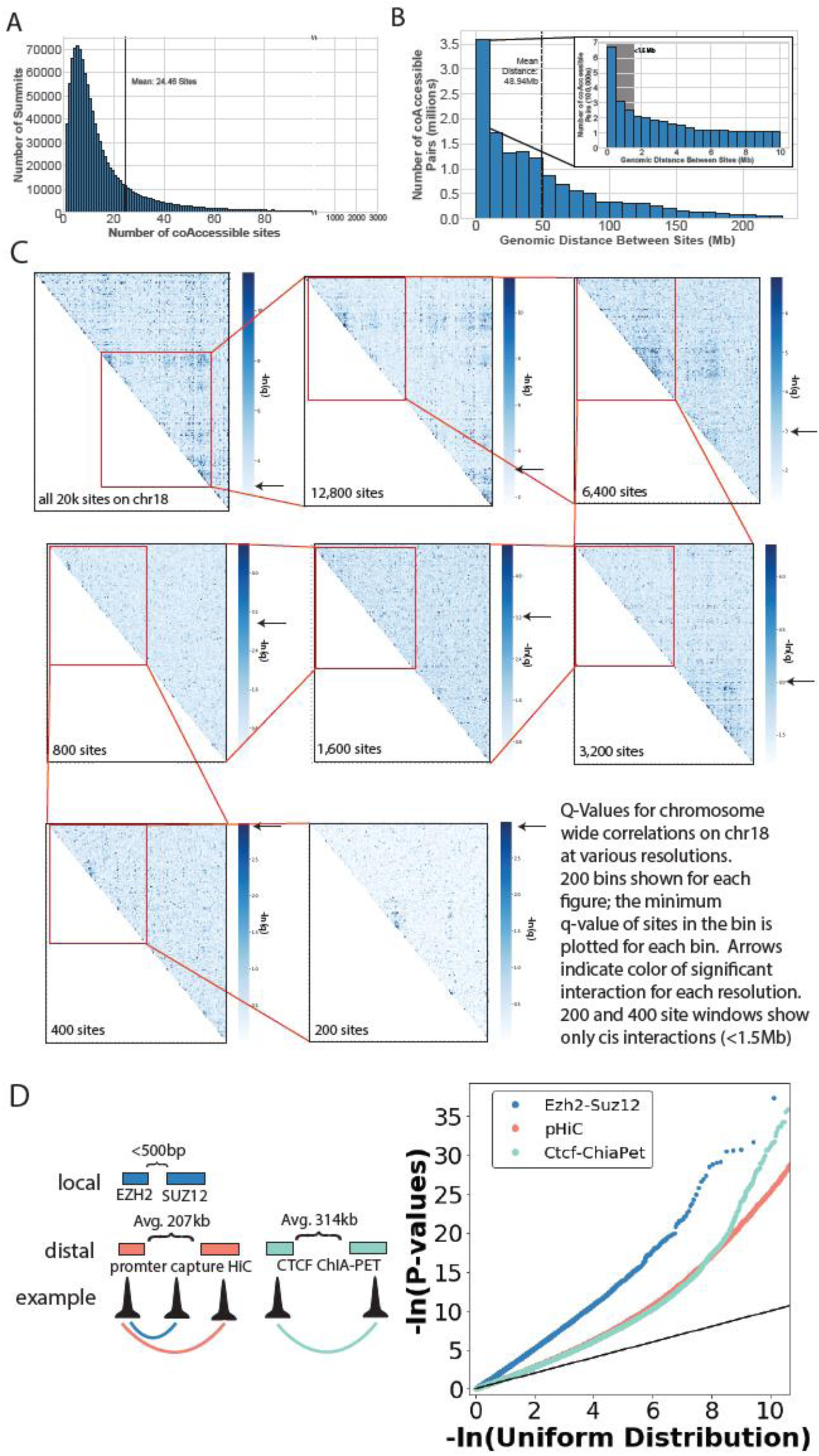
(A) Histogram showing the number of sites with a given number of co-accessible partners. The mean of 24.46 partners is highlighted by the dashed black line. (B) Histogram showing the distance between sites that are co-accessible. The mean of 48.94Mb is plotted with the dashed black line. The area highlighted in gray shows all co-accessible pairs at distance <1.5 Mb (ie previously studied distances). (C) Heatmap of regression q-values for various resolutions, ranging from the whole chromosome (top left) to 200 sites (bottom right), with 200×200 bins in each heatmap. Each box in red shows the region on the next zoom, starting in the top left and snaking to the bottom right. Each resolution is double the number of sites from the corresponding finer resolution. In the bottom right, each pixel in the heatmap is a single site (the grid is 200×200); in other panels, each pixel is the most significant q-value for all sites within the bin (ie for 400 sites, each pixel is 2-sites; for the entire chromosome, each pixel is ∼70 sites). Arrows on each color bar indicate color of a significant correlation (q = 0.05). At each resolution both local and distal significant associations can be seen. (D) QQ plot showing enrichments of p-values of co-accessibility combined across all chromosomes. (Left) Cartoon showing types of sites within each class: Ezh2-Suz12 TF pairs within 500bp of one another (blue), promoter capture HiC (red; avg 207kb apart) and CTCF ChIA-PET (teal; avg 314kb apart). Enrichments for local *cis* (blue) are above those for long-range *cis* (teal and red).

### Co-accessible sites are enriched for known biological processes

To determine whether calling co-accessibility across entire chromosomes reduced our power to identify *cis* co-accessibility, and/or resulted in technical artifacts, we examined the enrichment of co-accessibility for previously associated *cis* biological processes. First, using all co-accessible pairs from all chromosomes, we measured the enrichment for a local *cis* process: TF co-binding partners in the same protein complex (*EZH2* and *SUZ12* from PRC2^31,32^). We found that *EZH2*-*SUZ12* sites within 500bp of one another were highly enriched (Figure 2D, blue). Next, we examined enrichment for a long-range *cis* process: chromatin looping^6^. We found that opposing anchors of promoter centric chromatin loops measured via iPSC promoter capture HiC^33^ (mean 207kb apart; Figure 2D red), as well as structural chromatin loops measured via CTCF ChIA-PET^34^ from GM12878 (mean 314kb apart, Figure 2D teal), were also enriched. However, the enrichment for the long-range looping was lower than the local protein co-binding, consistent with *cis* effects being distance-dependent. Overall, these results show that the subset of co-accessibility within previously studied distances (<1.5Mb) recapitulates known *cis* characteristics of iPSCs.

### Co-accessibility intrinsically captures and predicts co-expression and chromatin state interactions

We performed unsupervised analyses to examine whether co-accessibility captures co-regulatory regions and genes, and if these relationships could be learned directly from co-accessibility. Since gene/protein regulation naturally has a network-like structure^35^, we modeled the co-accessibility as a network (Figure 3A). For each chromosome, we built an undirected graph with accessible sites as nodes, and edges between FDR q < 0.05 co-accessible sites weighted by their regression coefficient. We then annotated each site based on its overlap with gene promoters from GENCODE^36^, ROADMAP^12^ chromatin states for iPSC, and TF binding sites from ChIP-seq data in ESCs^31^ (see methods; Figure 3A). We first examined the correlation in gene expression (**co-expression**) for genes with co-accessible promoters using RNA seq data from the 154 iPSC lines^37^. We compared both the enrichment of co-expression correlation p-value (Figure 3B), as well as the proportion of tests that were significant (Figure 3C), across gene pairs that were stratified by distance up to 100Mb. As expected, we observed a high proportion (35%) of the examined *cis* gene pairs (<1.5Mb) whose promoters were co-accessible to be significantly co-expressed. Interestingly, we also observed a strong-enrichment for co-expression in pairs of genes with co-accessible promoters that were highly distal to one another (at least 10-100Mb apart, 27-30%). This enrichment was far greater than random distance matched genes (3%, Figure 3C, dashed line). Further, we found the proportion of significant tests to decay more quickly for genes within 1.5Mb of one another (slope = −0.26), compared to genes between 1.5Mb and 10Mb apart (slope = −0.03), suggesting the observed co-expression was largely driven by *trans* effects (as *cis*, but not *trans,* effects would be expected to decay with distance; Supplemental Figure 1C). These results reveal that co-accessible sites can *de novo* identify co-expressed gene pairs, and that while co-expression occurs most frequently in *cis*, it also occurs frequently across long ranges (including hundreds of megabases).

**Figure 3:**
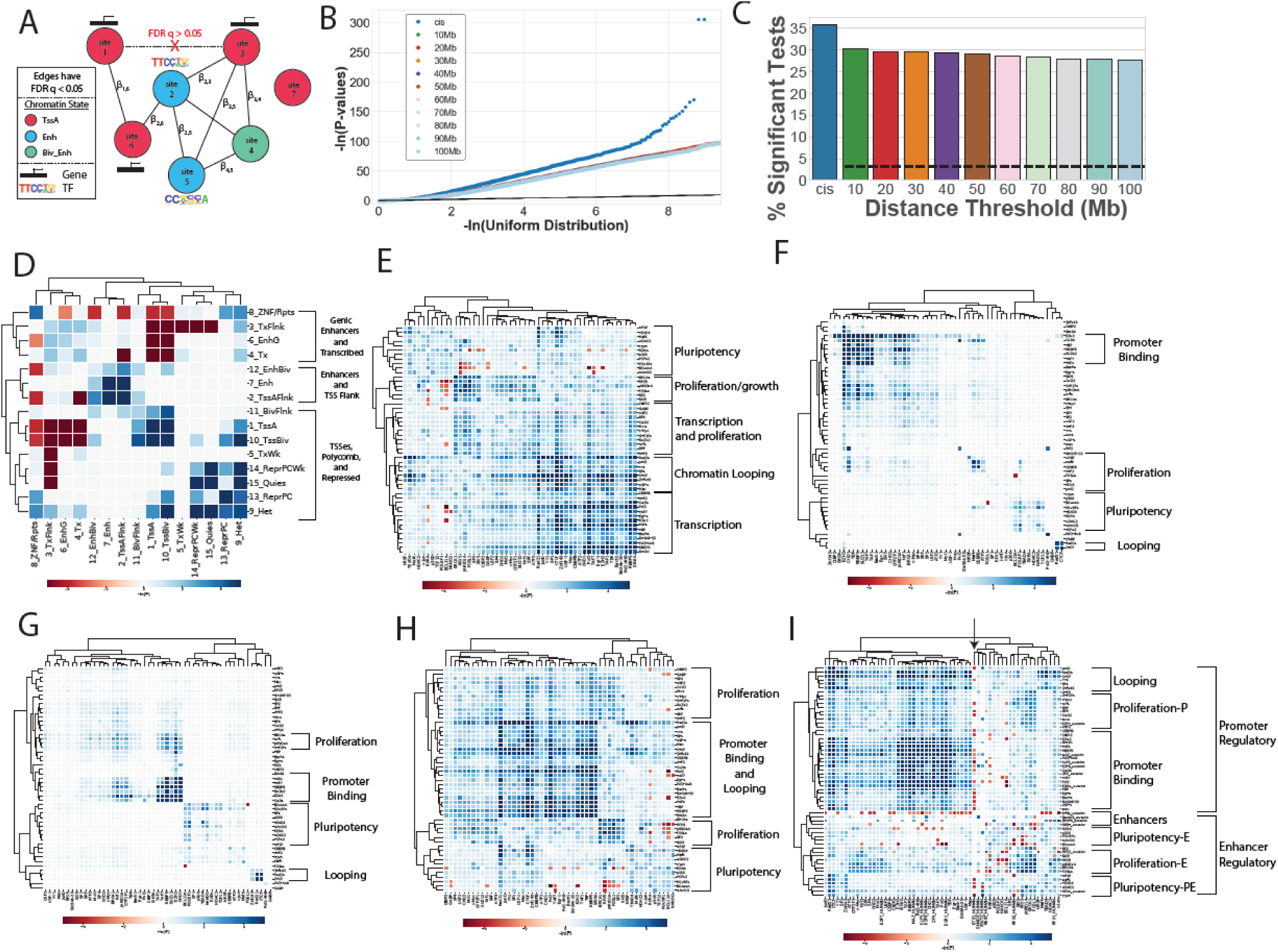
(A) Cartoon of a co-accessibility network. Accessible sites are represented as nodes, and FDR q < 0.05 co-accessible relationships are edges. For example, site 1 and 6 have a significant output from the LMM and thus have an edge; 1 and 3 do not. Edges are weighted by their regression coefficient (β). Nodes are labelled by their chromatin state (shown as colors), the genes whose TSS they overlap (shown as black boxes with arrows), and by the TF ChIP-seq peaks from ESC they overlap (shown as motifs). (B) QQ plot showing enrichments of p-values of correlation for co-expression of genes whose promoters are neighbors in the co-accessible network. Gene pairs are stratified by those that are at most 1.5Mb apart (*cis*) or those that are at least a given distance apart (colors; overlapping stratifications). *Cis* is enriched above all other distances, which are overlapping. (C) Bar plot showing the percent of co-expressed pairs that are FDR q < 0.05 at each distance threshold. Colors are shared between (B) and (C). (D-I) Heatmaps showing the signed empirical p-values of connectivity for (D) chromatin states, (E-H) TF ChIP-seqs, or (I) TF ChIP-seqs and predicted motifs. All distances (*cis* through 100Mb) shown for D, E, and I. (F) uses the local *cis* subnetwork induced from sites that are within 10kb of one another. (G) uses the long-range *cis* subnetwork induced from sites between 10kb and 1.5Mb apart. (H) uses the *distal* subnetwork induced from sites that are at least 1.5Mb apart. (D), (E), and (I) use chromosome 18. (F) and (G) use all edges from the genome-wide network. Clusters are labelled using the most common functionality of the included genes.

We next used the chromatin state annotations (Figure 3A) to examine co-accessibility between states (see methods). Due to computational constraints we focused on chromosome 18, observing three distinct clusters which highlighted known chromatin state interactions and iPSC specific biology^12,24,26-28^ (Figure 3D, Supplemental Figure 2 as a walkthrough): 1) genic enhancers and transcribed chromatin (active or weak; Supplemental Figure 2A); 2) enhancers, bivalent enhancers, and TSS flanking chromatin (Supplemental Figure 2B); and 3) active TSSes, bivalent TSSes, repressed/weak repressed, and heterochromatin (Supplemental Figure 2C). In addition to these 3 clusters, we found crossover between: 1) two different clusters (clusters 2 and 3) through Promoter-Promoter-Flanking interactions (Supplemental Figure 2D); and 2) two different subclusters (repressed and promoter in cluster 3) through active and bivalent TSS interactions with either strong or weak repressed polycomb (Supplemental Figure 2E). Overall, the observed gene co-expression and chromatin state clustering from unsupervised analyses suggest that co-accessibility can be used to *de novo* identify co-regulatory genes and chromatin states.

### Co-accessibility identifies novel co-regulatory TFs, as well as distance-dependent TF co-regulation

We sought to identify novel TF co-regulatory information captured by co-accessibility, and use it to derive new insights into the transcription factor landscape of iPSC gene regulation. Using the 51 ChIP-seq TF annotations on the co-accessibility networks (Figure 3A), we examined which pairs of TFs tended to be co-accessible more often than by chance (see methods). These transcription factor pair enrichments separated into five main clusters, (Figure 3E, Supplemental Figure 3 for a walkthrough) each of which contained numerous TFs that were known to be co-regulatory or functionally related: 1) pluripotency factors, including OCT4 (POU5F1), NANOG, and TEAD4 (Supplemental Figure 3A); 2) cell proliferation and organogenesis related TFs, including BRCA1, JARID1A, FOSL1, and SIX5 (Supplemental Figure 3B); 3) transcription and proliferation, including CtBP2, GABP, SP4, CHD2, and SRF for transcription, and c-Myc, AFT3, MXI1 and NRF1 for proliferation (Supplemental Figure 3C); 4) chromatin loop factors/structural factors, including Rad21, CTCF, YY1, SP1, JUND1, and Znf143 (Supplemental Figure 3D); and 5) transcription, including the promoter binding factors Pol2, TAF1&7, RBBP5, and TBP (Supplemental Figure 3E). While these five clusters recapitulated known TF groupings and functionality, we identified novel functions for TFs from cluster membership, subclustering, and cluster cross-over. As an example for cluster membership, RFX5 was a member of the proliferation/growth cluster (Supplemental Figure 3F), suggesting it may play a role in cancer; this is consistent with previous studies that found RFX5 upregulated in liver cancer, which results in the activation of genes associated with poor prognosis^38^. As a subclustering example, although Znf143 has been observed in promoter enhancer loops, it was not a member of the subcluster of the promoter enhancer specific loop TFs JunD, YY1, and SP1^39,40^ (Supplemental Figure 3G); rather, it had patterns similar to the broad loop factors Rad21 and CTCF, suggesting it may play a broad role in loop formation. As a crossover example, in the pluripotency cluster, we found some TFs (ex TEAD4, HDAC2) to have cross-over with the loop cluster, and others to not cross-over (ex BCL11A, NANOG; Supplemental Figure 3H), suggesting that some factors may have a more distal regulatory role than others. Overall, these analyses reveal that chromatin co-accessibility *de novo* recapitulates known gene regulation patterns and interacting TFs (including those that are cell type specific), and suggests that it may be possible to infer TF functionality from co-accessibility data.

As the examined networks contained co-accessible pairs at many different distances (from kilobases to megabases), we sought to find TF interactions unique to different regulatory distances. We stratified the network to interactions within 10kb (**local *cis***; Figure 3F), between 10kb and 1.5Mb (**long-range *cis***; Figure 3G), and greater than 1.5Mb (**distal**; Figure 3H). For the local and long-range *cis* networks, we used all chromosomes; for distal, we utilized chromosome 18 due to computational constraints. Across all three distance-stratified networks, we observed promoter binding, proliferation, pluripotency, and looping clusters; however, the specificity of these clusters, and the cross-overs between them, were different (Supplemental Figure 4 for walkthrough). For instance, in the local *cis* network, looping only contained CTCF and Rad21, whereas in the long-range *cis* network, the looping cluster also included Znf143, suggesting that Znf143 may act only on one anchor side for loop formation (Supplemental Figure 4A). Further, in both *cis* networks, the promoter cluster was separate from the loop cluster, consistent with only some of *cis* regulation involving chromatin looping; however, in the distal network, the promoter and loop clusters were combined, consistent with the majority of highly distal *cis* regulation utilizing chromatin loops. This suggests that the distal network contained *cis* interactions in addition to the interactions which span hundreds of megabases and are likely *trans* (Supplemental Figure 4A). We also observed stronger cross-over between the proliferation and promoter clusters in both the long-range *cis* and distal networks, compared to the local *cis* network (Supplemental Figure 4B), suggesting these TFs (ex JARID1A, GTF2F1) may mainly act through long-range and/or *trans* regulation. Finally, we observed more negative associations in the distal network than either *cis* network, suggesting that *trans* regulation may be comprised of more antagonistic regulation than *cis* regulation (Supplemental Figure 4C). Overall, these analyses suggest that TFs have different co-regulatory partners and directional relationships across different distances.

### Incorporation of motif predictions in identifying TF co-regulation reveals distinct promoter and enhancer clusters

To further examine what novel TF biology could be learned from co-accessibility, we expanded the network TF annotations to include predicted binding sites for TFs that were expressed and enriched in iPSCs (Figure 3I, Supplemental Figure 5 for a walkthrough). We identified two superclusters (Supplemental Figure 5A) which were composed of seven clusters that we named as follows: looping, promoter centric proliferation (proliferation-P), promoter binding, enhancers, promoter and enhancer centric pluripotency (pluripotency-PE), proliferation, and enhancer centric pluripotency (pluripotency-E). The two superclusters were separated by ETS1, with the top supercluster (containing Pol2) having a negative association. ETS1 has been shown to be a TF at sites occupied by Pol3 and involved in enhancer RNA transcription^41^. These observations suggest that TFs in the three clusters composing the supercluster anti-associated with ETS1 are primarily located at promoters (Supplemental Figure 5B), and TFs in the other supercluster (composed of four clusters; Supplemental Figure 5C) are primarily located at enhancers. Interestingly, we found proliferation clusters both in the promoter (ex TFs: NRF1, SRF) and the enhancer (ex TFs: JARID1, BRCA1) superclusters (Supplemental Figure 5D), as well as two different pluripotency clusters within the enhancer supercluster (ex TFs: OCT4, NANOG in pluripotency-E; TEAD4, HDAC2 in pluripotency-PE) (Supplemental Figure 5E). This suggests that TFs with similar functions may act across different genomic distances. While most of the promoter supercluster TFs were anti-associated with ETS1, and most of the enhancer supercluster TFs were not associated with ETS1, one cluster in the enhancer supercluster was anti-associated with ETS1 (Pluripotency-PE; Supplemental Figure 5F). This cluster (pluripotency-PE) also had a strong cross-over with the loop cluster (Supplemental Figure 5G), which in turn has a cross-over with the promoter binding cluster (Supplemental Figure 5H), suggesting that the TFs in the pluripotency-PE cluster regulate locally at promoters and distally through looping. Overall, these results show that co-accessibility can help delineate TFs that have primarily distal regulatory roles (Pluripotency-E and Proliferation-E clusters; Supplemental Figure 5I) from those that have primarily promoter regulatory roles (Proliferation-P and Promoter Binding clusters; Supplemental Figure 5J) from those which do both (Looping and Pluripotency-PE clusters; Supplemental Figure 5K).

### Identification of caQTLs and relation to co-accessibility

We sought to provide an in-depth characterization of how genetics is associated with total accessibility of sites (QTLs), as well as allele specific effects (ASE), in the context of *cis* and *trans* effects. We obtained genotypes for the 134 individuals from iPSCORE that had been previously identified using 50X WGS^37^, and tested for associations between the height of the accessible site and all genetic variants within 100kb^3,5^ of it using a linear mixed model. Across all chromosomes, we found 235k sites with an associated genetic variant within 100kb (***cis-* caSites**) with an FDR q < 0.05 (21%; Supplemental Table 2), which is consistent with previous estimates of the fraction of accessibility that is explained by variation^5^. We examined the enriched motifs at these *cis-*caSites, and found the top motifs enriched to be OCT4, CTCF, NANOG, SOX-family, and TEAD-family, consistent with iPSC gene regulation (Supplemental Table 3). We next examined the chromatin states enriched at these sites, and found an enrichment for non-promoter chromatin states (Figure 4A), suggesting that genetically associated sites were more likely to be distal regulatory in nature than located at gene TSSes or flanking chromatin. Next, we examined the association between co-accessibility and genetics by measuring the proportion of sites with a given connectivity that were significant *cis-*caSites (Figure 4B). We found that higher site connectivity corresponded to a higher proportion of significant *cis-*caSites, indicating that having more co-accessible partners increases the likelihood of having a *cis* genetic variant. This result suggests that having co-accessible partners allows for compensation against changes in a site due to genetic effects. Overall, these analyses identify sites whose total accessibility is genetically associated, and show that they are more likely to occur at iPSC distal-regulatory elements and to have high connectivity.

**Figure 4:**
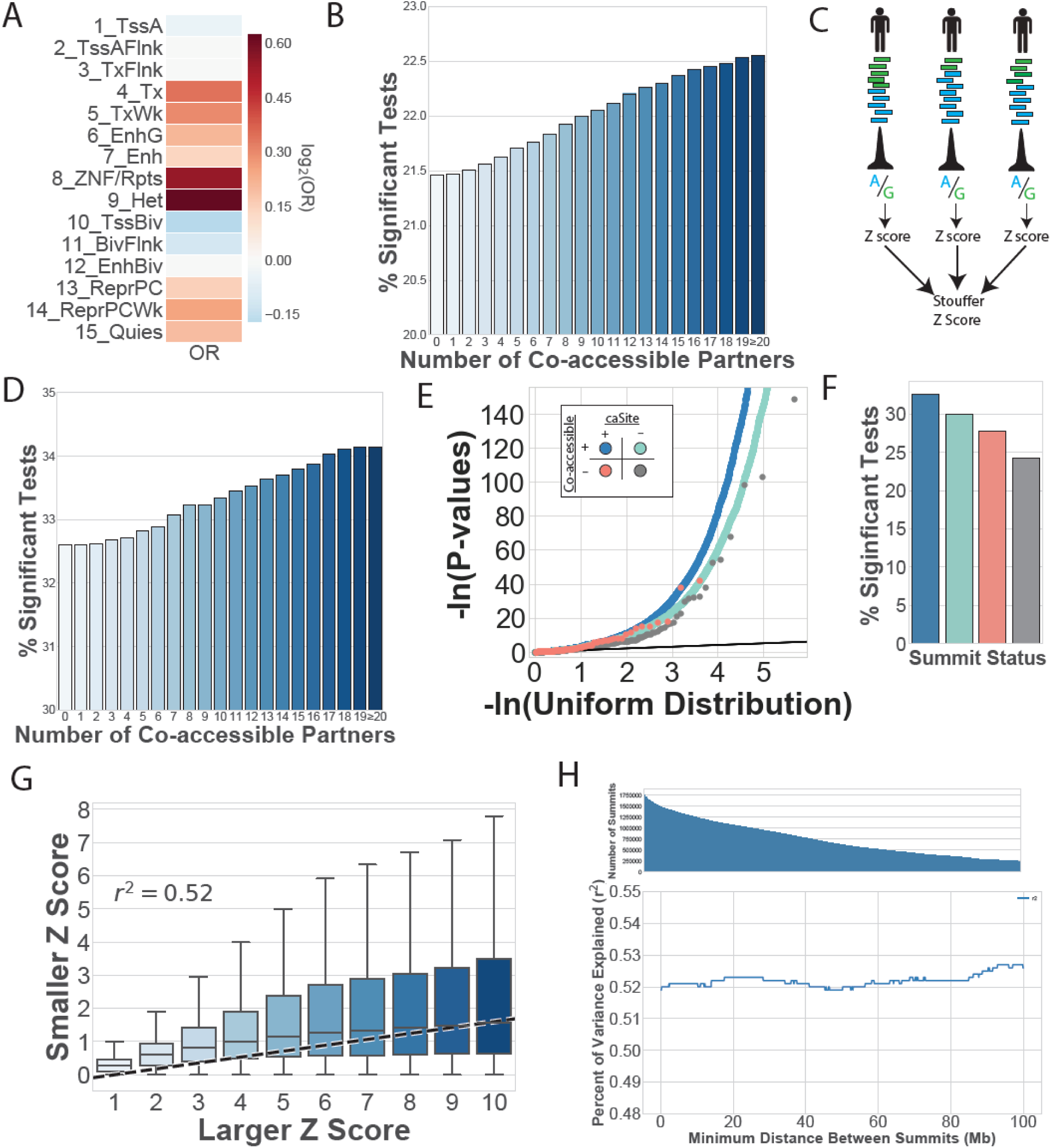
(A) Heatmap showing the log_2_ of the odds ratio of enrichment for chromatin states at *cis-*caSites compared to all accessible sites, with odds ratios set to 1 if the enrichment was non-significant. (B) Barplot showing the proportion of significant caSites as a function of the number of co-accessible partners in their networks. As sites have more co-accessible partners, they are more likely to have a *cis-*caQTL. Bars are colored by connectivity. (C) Workflow for identifying ASE for chromatin accessibility. Individual imbalance measurements were obtained per individual at sites with at least 10 reads. Z scores were calculated, and then combined across individuals for a single meta Z score per site. (D) Barplot showing the proportion of significant sites with ASE as a function of connectivity. As sites have more co-accessible partners, they are more likely to have exhibit ASE. Bars are colored by connectivity. QQ-plot and (F) Proportion of significant tests for ASE at sites that are co-accessible and caSites (blue), co-accessible and non-caSites (teal), singleton and caSites (red), or singleton and non-caSites (grey). Differences associated with caSites status can be seen by comparing blue to teal and red to grey. Differences associated with co-accessibility can be seen by comparing blue to red and teal to grey. (G) Boxplot of Z score relationship between co-accessible sites where both sites have heterozygous variants. Larger Z scores are plotted on the x-axis, and the paired smaller Z-score is on the y-axis. The regression line (calculated on the raw data) is plotted as a dashed black line. (H) Line plot showing the r^2^ value of larger ASE predicting co-accessible smaller ASE (as in panel G), restricting the analyses to sites at least X distance apart (up to 100Mb). The r^2^ holds consistent across all distances.

### Co-accessibility explains a large fraction of variation in ASE

To further probe the extent to which genetic effects were mediated by co-accessibility, we examined ASE. We measured ASE by calculating imbalance within each individual at all heterozygous variants within 200bp of an accessible site, and then meta-analyzing across individuals using Stouffer’s method (Figure 4C). This analysis identified >48,000 significant ASE sites at an FDR q < 0.05 (Supplemental Table 4) – notably, while ASE identifies regions associated with genetic effects, it does not delineate whether the imbalanced variant is causal for the effect, or neutral but in phase with a causal variant (ie proxy variant). We examined if sites with higher connectivity were more likely to exhibit ASE, and found that the proportion of significant ASE sites increased with connectivity (Figure 4D). This result suggests that co-accessibility may allow for a *cis* genetic effect to be mediated in *trans* to a co-accessible site (ie co-accessibility could be one of the processes which causes a proxy variant to be imbalanced).

To compare the relative effects of co-accessibility and *cis* genetic variation on ASE, we next we compared the distribution of ASE p-values (Figure 4E) and proportion of significant tests (Figure 4F) across four distinct sets of sites: 1) co-accessible *cis-*caSites (blue); 2) single *cis-*caSites (red); 3) co-accessible non-caSites (teal); and 4) single non-caSites (grey). As expected, we found both sets of caSites to be more enriched for ASE than their non-associated counterparts (Figure 4E,F blue vs teal, and red vs grey). Additionally, we found both sets of co-accessible sites to be more enriched for ASE than non-co-accessible sites (Figure 4E,F, blue vs red, and teal vs grey), consistent with Figure 4D. Interestingly, we found co-accessibility status to be more enriched for ASE than *cis-*caSite status (Figure 4F, both blue and teal are enriched above red). This result further supports *trans* genetic effects being mediated through co-accessibility, as these sites do not have a significant *cis-*caQTL, but do show significant allelic effects. To make sure that this observation was not predominantly driven by false negative caQTLs, we examined whether ASE in one site could explain ASE in co-accessible sites by regressing the lead Z score of a co-accessible network against each partner Z score en masse across all chromosomes simultaneously (see methods). We found a large fraction of variation in ASE to be explained by the single most imbalanced co-accessible site (r^2^=0.52, p < 10^-32^; Figure 4G), showing that genetic variants can exert *trans* effects via co-accessibility. Finally, we examined whether this high predictability was consistent across varying genomic distances. We found the r^2^ to hold surprisingly constant up to 100Mb apart, ranging between 0.51 and 0.53 (Figure 4H), despite a large difference in the number of pairs used in the model (between 250k and 1.25M). Together, these results suggest that an accessible site can be influenced in *trans* by a distal genetic variant through intermediate effects on a co-accessible partner.

### Identification of trans-caQTLs by leveraging co-accessibility network

We set out to identify genetic variants that indirectly affect distal sites by mediating their *cis* effects through co-accessibility (ie *trans-*caQTLs). To identify *trans-*caQTLs within the same chromosome, we leveraged the co-accessibility network to perform targeted association tests, thereby reducing multiple hypothesis testing. To perform these analyses, we restricted our tests to variants that were *cis-*caQTLs, and tested them against neighbors of the respective *cis-*caSite in the co-accessibility network that were at least 1.5Mb away (Figure 5A). This identified 368,639 putative *trans-*caQTL-caSite pairs out of a tested 9,967,402 pairs (3.7%) at an FDR q < 0.05 (Supplemental Table 5). Notably, many of these putative *trans-*caQTLs were highly distal to their targets (Figure 5B), with some hundreds of megabases away. However, this regression analysis cannot delineate a true *trans* interaction in which the variant’s effect on an accessible site is mediated through a co-accessible partner (Figure 5C left) from two independent *cis* effects driven by the same variant (Figure 5C right). We thus further probed these putative *trans*-caQTLs by performing a mediator analysis to identify variants with a statistically significant fraction of their association with the *trans*-caSite explained by the height of the *cis*-caSite^42^. For these analyses, we tested all genotypes in the 134 individuals (ie if the site was multiallelic, we included all alleles; 80% of multiallelic sites were indels). Out of the 934,136 putative *trans*-caQTL genotypes, the mediator analysis found 92,638 to be significantly mediated at an FDR q < 0.05 (9.9% of the putative, ∼1% of all tests; Supplemental Table 6). As sites have high connectivity, it is possible that a given cis-caSite which mediates a *trans* effect (*cis-*mediator-caSite) exerts its effects on multiple co-accessible partners. We thus examined whether *cis-* mediator-caSites affected multiple co-accessible *trans-*caSites, and found the 92,638 *trans*-caSites to be mediated by 29,362 *cis-*mediator-caSites, with a mean of 3.16 *trans-*caSites per mediator (Figure 5D). This suggests that when a variant affects an accessible site, the effects are mediated throughout its co-accessibility network, rather than in a pair-wise fashion with only one of the co-accessible partners. We compared the motifs underlying *cis-*mediator*-*caSites and *trans-*caSites, and found both to be similarly enriched for OCT4, CTCF, NANOG, SOX-family, and TEAD-family motifs (Supplemental Table 3). We next examined the chromatin states at *cis-* mediator-caSites and *trans*-caSites, and found both to be enriched for promoter and gene centric chromatin states (Fisher’s Exact FDR q < 0.05; Figure 5E); however, only *trans-*caSites were depleted at enhancers and bivalent enhancers. The fact that these enrichments are different suggests that genetic effects have a directionality within co-accessibility networks. Together, these analyses identified tens of thousands of *trans-*caQTLs, suggests that *trans* effects are directionally mediated throughout a network, and shows that co-accessibility can be leveraged to identify *trans*-caQTLs from a relatively small sample size.

**Figure 5:**
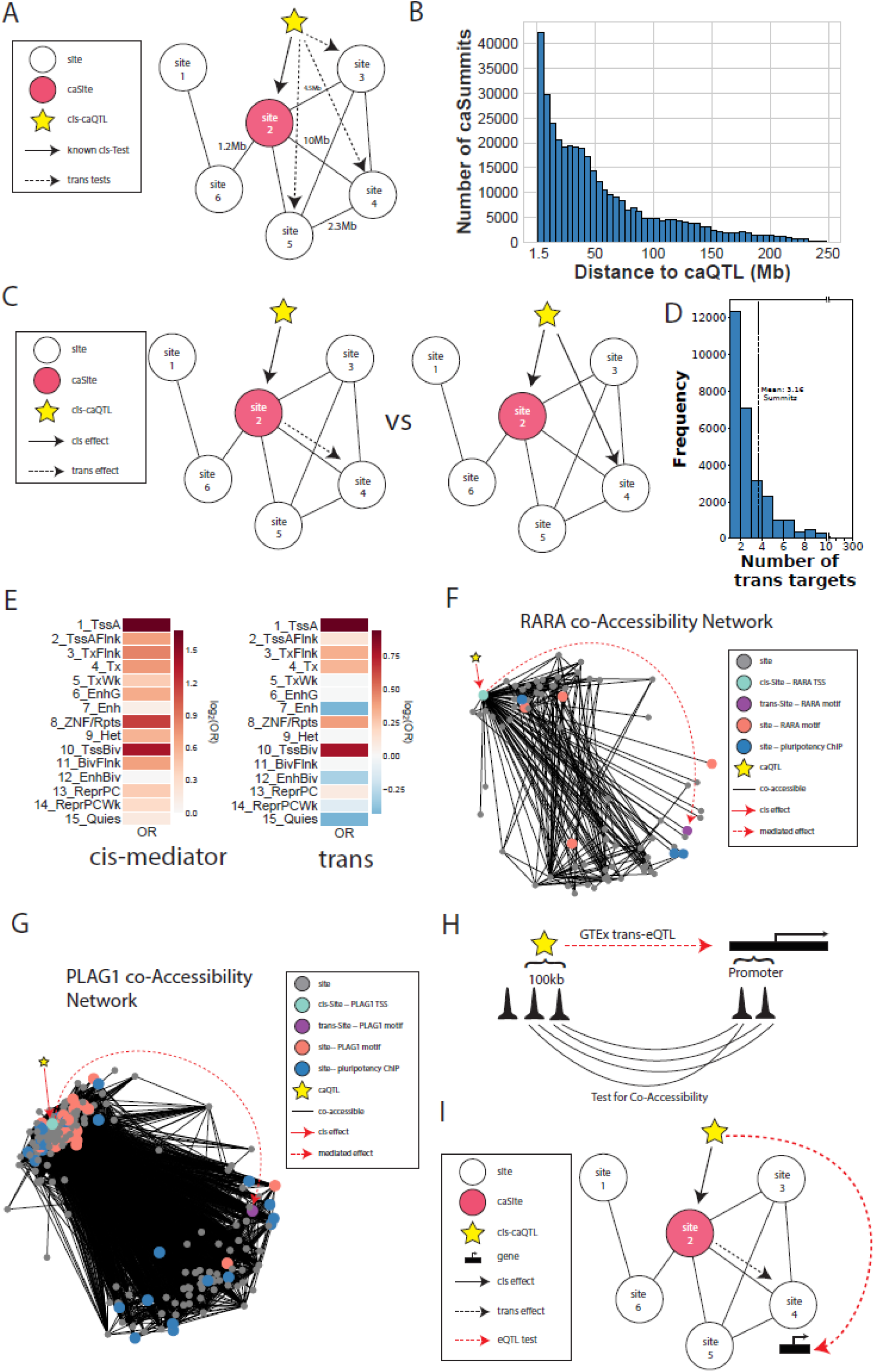
(A) Cartoon illustrating how putative *trans-*caQTLs were tested. *Cis*-caQTLs (star) were tested as *trans*-caQTLs against the neighbors (sites 3,4, and 5) of the *cis-*caSite (site 2) in the co-accessibility network. (B) Histogram showing the distance distribution of significant putative *trans-*caQTLs to their target *trans-*caSite. (C) Cartoon illustrating mediator analysis, which tests whether a variant exerts a *trans* effect on a site (site 4) through an intermediate site (site 2; left), or whether the variant exerts two independent effects (right). (D) Distribution of the number of mediated *trans-*caSites per *cis-*mediator-caSite. (E) Heatmaps showing the log_2_ of the odds ratio of enrichment for chromatin states at (left) *cis-*mediator-caSites or (right) *trans-*caSites compared to all *cis-*caSites, with odds ratios set to 1 if the enrichment was non-significant. (F and G) Co-accessibility networks centered on two particular sites: (F) an accessible site at the RARA promoter, and (G) an accessible site at the PLAG1 promoter. Nodes are accessible sites, edges show significant co-accessibility. Node color indicates whether the node is the *cis-* medaitor-caSites which defines the network (teal), the *trans*-caSite containing the TF for the respective gene (purple), a non-associated site containing the respective TF (red), a site containing an iPSC ChIP-seq factor (blue), or a different site (grey). The caQTL is shown as the star, with a solid red line for its *cis*-effect, and a dashed red line showing the mediated effect from the *cis-*mediator-caSite to the *trans*-caSite. (H) Cartoon illustrating tests for co-accessibility at GTEx *trans-*eQTLs. Sites within 100kb of the *trans*-eQTL variant were tested against sites at the eGene promoter. (I) Cartoon illustrating how *trans* eQTLs were tested. Sites with a significant mediator q value (site 2) had their *cis-*caQTLs (star) tested as an eQTL against the genes which had co-accessible promoters (site 4).

To gain better insight into the mechanisms underlying these *trans-*caQTLs, we characterized two large co-accessibility networks centered on *cis-*mediator-caSites. These networks were chosen because their *cis*-mediator-caSite was at the promoter of a TF, and their *trans-*caSite contained a binding site for that TF. The smaller of these two networks was on chromosome 17 (Figure 5F) with 77 total sites, 30 of which overlapped gene promoters. One of these 30 sites was at the RARA gene promoter, and was a *cis-*mediator-caSite whose *trans-*caSite overlapped a RARA binding motif. Four other sites were also at RARA binding sites. The RARA gene has been implicated in development, differentiation, and transcription of clock genes^43^; we therefore examined the network for iPSC TFs, and found three sites overlapping ChIP-seq binding sites for the core pluripotency TFs (OCT4, NANOG, and TEAD4). Finally, we examined the function of the 30 genes whose promoters were in the RARA network, and found their proteins to be statistically enriched for having protein-protein interactions (**PPIs**) in StringDB (StringDB enrichment p = 9.65×10^-3^; Supplemental Figure 6A). Further, the functionality of these genes was enriched for gene sets for multiple cancer types, including acute myeloid leukemia (AML) and Breast Cancer (Supplemental Figure 6A), suggesting co-accessibility network dysregulation may play role in cell-type relevant disease. The second example, on chromosome 18, was a co-accessibility network comprised of 261 sites, of which 130 were at gene promoters (Figure 5G). One of these 130 sites was at the PLAG1 promoter, and was a *cis*-mediator-caSite for a *trans-* caSite at a PLAG1 motif. In addition to the *trans-*caSite, 23 other sites contained PLAG1 motifs. PLAG1 is developmentally regulated^43^; we therefore also examined iPSC TFs, and found 24 sites overlapping a ChIP-seq peak for NANOG, OCT4, or TEAD4. Finally, the proteins transcribed by the genes in the network were significantly enriched for being having PPIs (StringDB enrichment p = 8.12×10^-4^; Supplemental Figure 6B), but not for any StringDB gene sets. Notably, this set proteins contained 19 experimentally validated PPIs. Overall, these analyses show that co-accessibility can be used to identify novel *trans* regulatory modules which can be disease-associated.

**Figure 6:**
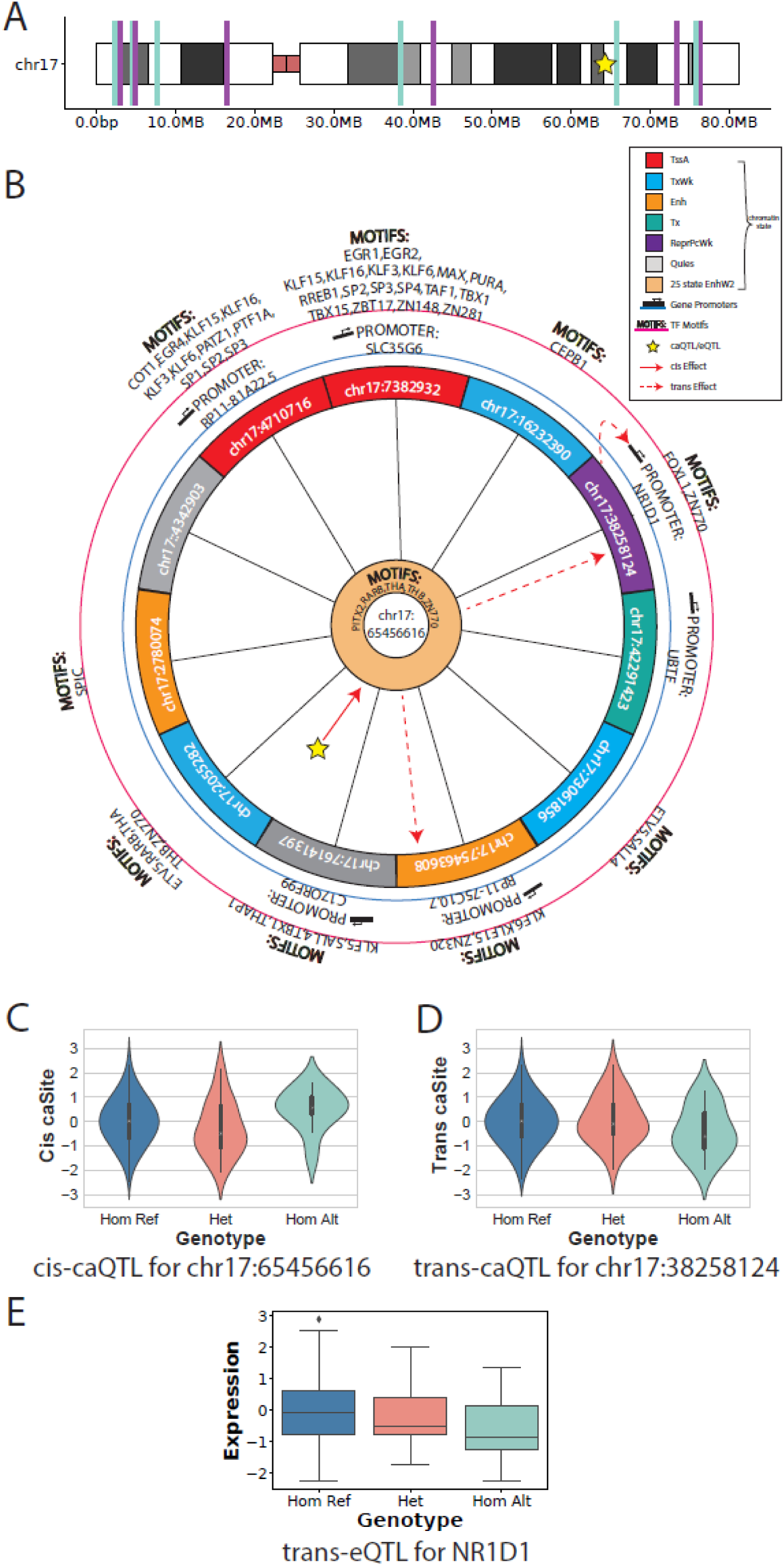
An overview of the chr17:65456616 co-accessible network. (Top) ideogram showing location of sites across the chromosome. Two colors are used so that nearby sites are visible. The star indicates the caQTL/eQTL. (Middle) The co-accessibility network; nodes shown as the inner circle (chr17:6546616) or partitions of the outer circle (neighbors). Black lines (spokes) are co-accessible relationships, solid red line shows *cis-*QTL effect, and dashed red lines show mediated *trans-*caQTL effects. Colors on the partition show the chromatin state, the blue circle lists gene promoters overlapped, and the red outer circle lists Motifs that are overlapped. (Bottom) Violin or boxplots of QTL results for the *cis*-medaitor-caQTL for crh17:65456616, the *trans-*caQTL for chr17:38258124, and the *trans-* eQTL for NR1D1. All effects are from the same variant.

### Identification of *trans*-eQTLs from *trans*-caQTLs at promoters

It is possible that some of variants affecting chromatin accessibility could propagate their effects to changes in gene expression. We hypothesized that some of the variants underlying *trans*-caQTLs whose *trans-*caSite was at a promoter for a gene were also *trans*-eQTLs for that gene. As previous studies have shown that a large cohort is required for sufficient power to detect *trans*-eQTLs^4^, we initially examined whether *trans-*eQTLs previously identified in GTEx (in different tissues and inter-chromosomally) exhibited co-accessibility with their eGene in our data (Figure 5H). We performed a targeted inter-chromosomal analysis (as our networks were all intra-chromosomal), examining the 32 non-MHC *trans-*eQTLs in GTEx for genes expressed in the iPSCORE iPSCs. Surprisingly, we found that 97% (31/32) of these *trans*-eQTLs had co-accessibility between a site at the promoter of the eGene and one near the eQTL (FDR q< 0.05, see methods), despite none of these *trans*-eQTLs being discovered in stem cells. This result suggests the majority of *trans*-eQTLs have co-accessibility associated with them, and that co-accessibility may be conserved across cell types.

Next, we sought to determine whether *trans-*caQTLs could inform and increase power for detecting *trans-*eQTLs. We performed an intra-chromosomal *trans-*eQTL by identifying all genes whose promoters overlapped a *trans*-caSite, and used an LMM to test for association between gene expression and the genotype of the *trans*-caQTL (Figure 5I). Overall, we found an enrichment within the *trans-*eQTL p-value distribution (λgc=1.49, Supplemental Figure 6C), and 9 significant *trans-*eQTLs (FDR q < 0.05; Table 1). These results show that co-accessibility can be utilized to increase power in detection of *trans-*eQTLs, as GTEx (a larger multi-tissue study) identified an average of 2.7 *trans-*eQTLs per tissue. The 9 *trans*-eGenes were *RASSF7, EHMT1, DPP9, LMNB2, RGS3, AC009133*.*17, SDCCAG8, PDE2A*, and *NR1D1,* which are all related to iPSC functionality (ie cell cycle, growth, division) or relevant diseases (ie cancer)^43^. The median distance between the corresponding eQTL and eGene was 27Mb; one pair was over 200Mb apart. Within the average 27Mb window of a gene, the 134 individuals in this study had ∼400k variants; thus, our approach of only testing *trans-*caQTLs against expression levels of genes with *trans*-caSites at their promoters greatly reduced the p-value threshold for significance. These results demonstrate the advantages of using co-accessibility to identify *trans* effects on cell type specific gene regulation.

### Integrating co-accessible annotations to infer trans-eQTL mechanisms

Finally, to characterize how chromatin accessibility, gene expression, regulatory variation, chromatin states, TFs, connectivity, and genomic distance fit within the context of co-accessibility, we visualized one of the networks from the *trans*-eQTL analysis (Figure 6). The network centered on the *cis-*mediator-caSummit (chr17:65456616) contains 12 co-accessible sites, spanning both the P and Q arms of chr17 (Figure 6A). The *cis-*caQTL for the *cis-*mediator-caSummit (chr17:65456616; Figure 6C) is mediated to two *trans-*caSites, chr17:38258124 and chr17:75463608 (Figure 6B, dashed red lines; Figure 6D). One of these *trans*-caSites, chr17:38258124, is at the *NR1D1* promoter (Figure 6B) which is associated with circadian rhythm and reported to have iPSC specific functionality^44^. The *cis-*caQTL is also a *trans-*eQTL for *NR1D1* (Figure 6E). In this network, the 11 sites co-accessible with *cis-*mediator-caSummit chr17:65456616 were not co-accessible with one another (Figure 6B; the hub and spoke shape of the network). These 11 sites are at 6 different types of chromatin states, 6 gene promoters, and contain numerous TF motifs (Figure 6B); and the *cis-*mediator-caSite is at an iPSC weak enhancer (EnhW2 from the 25-state model of E020 in ROADMAP) and contains motifs for PITX2A, RARB, THA, THB, and ZN770. Together, these data suggest that the *trans-*eQTL, which is 27Mb distal from its eGene *NR1D1*, exerts its effects by modulating the binding of one or more of the 5 TFs at the *cis-*mediator-caSite. Overall, these data exemplify how annotating co-accessibility networks with multiple types of molecular phenotypes can identify *trans* genetic effects and putative mechanisms underlying them.

## Discussion

Here, we performed ATAC-seq in 152 iPSC lines from 134 individuals, and use the data to find chromosome-wide co-accessibility. We show co-accessibility is highly connected, with sites being co-accessible with an average of 24 other sites, and can span long distances (up to hundreds of megabases). We then show that, from annotated co-accessibility alone, it is possible to find, *de novo,* co-regulatory chromatin states, genes, and TFs. Additionally, we use this information to infer novel TF functionality and observe that binding sites for TFs with similar functions or that act in complexes are co-accessible at long distances (up to hundreds of megabases). Finally, we perform one of the largest chromatin accessibility QTLS (caQTLs) to date, identifying hundreds of thousands of *cis-*caQTLs, tens of thousands of *trans-*caQTLs, and 9 *trans-eQTLs* as well as putative mechanisms underlying them.

We show that chromatin co-accessibility is a mechanism by which distal *trans* genetic effects are mediated. We found that co-accessible sites were more likely to have a *cis* genetic effect, and that allelic effects were predictive of co-accessible allelic effects. Additionally, we also show that genetic variants that are associated with accessibility often mediate their effects to multiple distal partners through co-accessibility. Together, these results suggest that co-accessibility may function as an insulator, allowing a given regulatory system to be more robust to perturbation by having sites compensate for their co-accessible partners. Future studies examining the effects of perturbing multiple aspects of the same co-accessibility network in a dose dependent manner could validate this hypothesis, as well as provide insight into the spreading of *cis* genetic effects to *trans* throughout the genome.

Previous studies^4^ utilizing large cohorts of individuals and multiple tissue types have shown that it is difficult to properly power a study for the identification of *trans-*eQTLs. We show that co-accessibility data can be practically used to reduce the multiple testing burden faced by genetic association studies due to the large search space for *trans* effects. By leveraging co-accessible information, we were able to test single variants against single sites, enabling the identification of tens of thousands of *trans*-caQTLs from only 134 individuals. Further, these analyses translated to gene expression, with 9 *trans-*caQTLs also being *trans-*eQTLs for iPSC relevant genes. These 9 eGenes were identified only examining intra-chromosomal *trans* effects, and only in one tissue, compared to GTEx which had 94 *trans* eGenes across ∼50 tissues. Future studies could perform ATAC-seq and RNA-seq in the same individuals to define co-accessibility networks, and then use them to direct the identification *trans*-eQTLs to increase statistical power. Despite our high computational power (a 16 node, 512 core compute cluster with 2.25TB of RAM total), we were only able to test for intra-chromosomal co-accessibility due to computational requirements (this process took multiple months of running LMMs) and a relatively small cohort – yet found tens of thousands of *trans-*caQTLs. Future studies with more power and resources could likely utilize inter-chromosomal co-accessibility to define co-accessibility networks across all pairs of chromosomes and enable the identification of even more *trans*-caQTLs and *trans-*eQTLs.

## Contact Information

Kelly Frazer:

## Acknowledgements

This work was supported in part by a California Institute for Regenerative Medicine (CIRM) grant GC1R-06673 and NIH grants HG008118-01, HL107442-05, DK105541-03 and DK112155-01. RNA-seq were performed at the UCSD IGM Genomics Center with support from NIH grant P30CA023100. WWYG was supported by the National Heart, Lung, and Blood Institute of the National Institutes of Health under Award Number F31HL142151. Whole genome sequencing was performed at Human Longevity, Inc.

## Author Contributions

Conceptualization, A.C.D, W.W.Y.G., and K.A.F.; Methodology, A.C.D., W.W.Y.G., and E.N.S; Software, W.W.Y.G.; Validation, W.W.Y.G.; Formal Analysis, W.W.Y.G.; Investigation, A.C.D., W.W.Y.G., P.B.; Data Curation, A.C.D., W.W.Y.G., H.M., M.D.; Writing – Original Draft, A.C.D., W.W.Y.G., M.D., and K.A.F.; Visualization, W.W.Y.G.; Supervision, A.C.D, and K.A.F.; Project Administration, A.C.D., and K.A.F.; Funding Acquisition K.A.F.

## Supplemental Figure Legends

**Supplemental Figure 1:** (A) Barplot showing the number of edges (blue) in the co-accessibility network (ie co-accessible pairs), or number of edges normalized to chromosome size (red). (B) Scatter plot showing the regression βs from co-accessibility LMMs for significant results, plotted against the linear distance between sites on the genome. Red lines indicate the average β for each 1Mb bin – the reduction of individual high absolute βs as distance increases, but consistent average, indicates strength does not decay with distance, but the number of co-accessible pairs does. (C) Barplot showing the proportion of significant co-expressed pairs that are at most X distance apart from one another. The plot elbows off near 1.5Mb (from visual inspection); the slopes are shown for bins <1.5Mb (green), and from 1.5Mb to 10Mb (red).

**Supplemental Figure 2:** A walkthrough of the chromatin-state co-accessibility unsupervised clustering analysis. The same heatmap is shown in each panel, with specific points highlighted for each panel: (A) The genic and transcribed cluster; (B) the Enhancer and TSS Flank cluster; (C) The TSSes, Polycomb, and Repressed cluster; (D) Crossover between the Enhancer and TSS Flank cluster with the TSSes, Polycomb, and Repressed cluster via TssA_Flnk-TSS association; (E) Crossover between subclusters in the TSSes, Polycomb, and Repressed cluster.

**Supplemental Figure 3:** A walkthrough of the TF co-accessibility unsupervised clustering analysis. The same heatmap is shown in each panel, with specific points highlighted for each panel: (A) The pluripotency cluster and the specific TF members mentioned in the text; (B) The proliferation/growth cluster and the specific TF members mentioned in the text; (C) The transcription/proliferation cluster and the specific TF members mentioned in the text; (D) The chromatin looping cluster and the specific TF members mentioned in the text; (E) The transcription cluster and the specific TF members mentioned in the text; (F) Inferring RFX5 functionality from membership within the proliferation/growth cluster; (G) Inferring Znf143 functionality from subclustering within the loop cluster; (H) Inferring distal functionality from specific TFs in the pluripotency cluster from crossover between these TFs and the chromatin looping cluster.

**Supplemental Figure 4:** A walkthrough of the distance stratified TF co-accessibility unsupervised clustering analysis. The same 3 heatmaps are shown in each panel, with specific points highlighted for each panel: (A) The looping cluster contains CTCF and Rad21 for the local *cis* network, adds Znf143 in the long-range *cis* network, and is combined with the promoter binding cluster in the distal network; (B) Crossover between the pluripotency and promoter cluster increases from local *cis* to long-range *cis* to distal networks, with the two TFs mentioned in the text (GTF2F1 and JARID1A) not involved in local *cis,* clustered together in long-range *cis,* and crossing over in *distal*; (C) Comparison of anti-associations found across networks.

**Supplemental Figure 5:** A walkthrough of the TF ChIP and Motif co-accessibility unsupervised clustering analysis. The same heatmap is shown in each panel, with specific points highlighted for each panel: (A) the two superclusters of the network (Promoter Regulatory and Enhancer Regulatory); (B) the three clusters comprising the Promoter Regulatory supercluster; (C) The four clusters composing the Enhancer Regulatory supercluster; (D) Both proliferation clusters observed; (E) both pluripotency clusters observed; (F) The pluripotency-PE cluster; (G) Crossover between the pluripotency-PE cluster and the looping cluster; (H) Crossover between the looping cluster and the promoter binding cluster; (I) Clusters (Pluripotency-E and Proliferation-E) that are primarily enhancer associated, with example TFs mentioned in the text (NANOG, JARID1A, BRCA1) highlighted with arrows; (J) Cluster (Proliferation-P) that is primarily promoter associated with example TFs mentioned in the text (Nrf1 and CHD2) highlighted with arrows; (K) Cluster (Pluripotency-PE) that associates with both promoters and enhancers, with example TF mentioned in the text (TEAD4) highlighted with an arrow.

**Supplemental Figure 6:** (A and B) StringDB results from the genes whose promoters are in the (A) RARA network and (B) PLAG1 network. Protein network (top in both pannels) and Top GO enrichments in reference publications (bottom in A) are shown. Colors for edge connections in the protein networks come from StringDB and are known from (teal) curated databases or (pink) experimentally validated; predicted from (green) gene neighborhoods, (red) gene fusions, or (blue) gene co-occurrence; or associated together via (light green) text mining, (black) co-expression, or (light blue) protein homology. (C) QQ-plot for *trans-*eQTL, with zoom between 0 and 1 on both axes shown as a subplot with λ_gc_ drawn and labelled.

**Supplemental Figure 7:** QC and filtering results for ATAC peaks using the more precise sub-peak structure within them (ie summits). (A) Distribution of TMMs at summits (B-D) Statistics and enrichments after filtering out peaks where no summit had a median coverage across individuals (**IndivMed**) greater than or equal to 0, 1x the median, or 2x the median of the IndivMed across all summits (**SummitMed**). For example, TMM threshold 2 contains all peaks where max(IndivMed) ≥ SummitMed × 2. (B) Barplot showing number of peaks in each set. (C) TMM distribution of filtered peaks (Teal) compared to original peaks (Blue). (D) Chromatin state enrichments. (E) Distribution of peak size of the 1-TMM filtered set (ie the set used throughout the manuscript), with the median peak size shown via the dashed line.

## Supplemental Table Legends

Supplementary Table 1: Individuals and Data Used in this study

Supplementary Table 2: Significant caSummits and Lead Variants

Supplementary Table 3: HOMER Motif Enrichment Results for cis-caSites, cis-mediator-caSites, and trans-caSites

Supplementary Table 4: ASE Meta-analysis Results

Supplementary Table 5: Significant Putative Trans eQTL Results

Supplementary Table 6: Significant Trans-caQTL Mediator Results

## Methods

### Selection of Individuals form iPSCORE

152 iPSC lines from 134 individuals from iPSCORE were selected for ATAC-seq analysis. These 134 individuals are from multiple ethnicities; among them, 82 individuals belong to 26 families and 52 are unrelated. For all 152 lines, ATAC libraries were generated from matched iPSC and iPSC-derived cardiomyocytes (cardiomyocytes not part of this manuscript).

### ATAC-seq

We performed ATAC-seq on 152 iPSC samples using the protocol from Buenrostro et al. (Buenrostro et al., 2013) with small modifications. Frozen nuclear pellets of 2.5 × 10^4^ PSCs were thawed on ice and tagmented in total volume of 25μl in permeabilization buffer containing digitonin and 2.5μl of Tn5 from Nextera DNA Library Preparation Kit (Illumina) for 45-75min at 37°C in a thermomixer (500 RPM shaking). To eliminate confounding effects due to index hopping, all libraries within a pool were indexed with unique i7 and i5 barcodes. Libraries were amplified for 12 cycles using NEBNext® High-Fidelity 2X PCR Master Mix (NEB) in total volume of 25µl in the presence of 800nM of barcoded primers (400nM each) from custom synthesized by Integrated DNA Technologies (IDT). Each library was independently sequenced twice on Illumina HiSeq 4000 with paired-end 150bp reads.

### ATAC Peak Calling

Peaks were called using MACS2 v2.1.1.20160309^45^ with the settings: --nomodel --nolambda -- keep-dup all -f BAMPE -g hs. Peaks were called either individually, or simultaneously on all samples by providing each input sample to MACS2 at the same time with the -t option.

### Identification and QC of ATAC-seq peaks

To assess the quality of each sample, we identified and characterized peaks per sample. We first aligned the two sequencing runs for each of the 152 samples individually (304 BAM files), removed duplicates, and to ensure that identified peaks represented TF binding sites rather than Tn5 insert sites flanking nucleosomes, filtered to read inserts ≤140bp in length. Following this processing, we separately called peaks on each of these 304 BAM files using MACS2. To assess the quality of each sample, we examined the fraction of reads in peaks and the percent of peaks falling within active regions of the genome as defined by ROADMAP chromatin states 1,2,3,5,6,7, and 11 for iPSC (E020). We found the mean FRiP of the 304 samples to be 13%, and the mean percent of peaks in active regions to be 50%. We merged the two fastq files for each sample, re-removed duplicates, and re-filtered to read inserts ≤140bp in length, producing the final set of 152 BAM files for downstream analyses.

### Determining sample coverages

Coverages were obtained using the featureCount package from subread v1.5.0^46^ for each sample individually on a set of peaks or summits. Next, counts were TMM-normalized using edgeR v3.12.1^29^ for each peak call or summit call set across all individuals.

### Creating a reference set of ATAC-seq summits across the 152 samples

ATAC-seq identifies punctate regions of accessible chromatin which demarcate transcription factor (**TF**) binding sites. However, as peaks must be called from the data *de novo*, rather than identified from a set of known sites as per gene expression analyses, it is important to identify a consistent “reference” set of ATAC-peaks that can be compared consistently across samples. We thus called peaks and summits using MACS2 on all samples simultaneously, identifying a total of 859,563 peaks with 1,839,425 summits. We sought to filter out broad regions with low coverage while maintaining peaks that potentially contained multiple real TF binding sites. We started by finding the normalized coverage in TMMs for each sample across all ∼1.8M summits. We found the vast majority of summits to have a median of ≤ 2.0 TMMs across the 152 samples (Supplemental Figure 7A). As the summits represent the high points in the peaks, we filtered the peaks based on the median coverage of the contained summits (medCov). Specifically, we tested three filters based on the maximum medCov of all summits within each peak: maximum median coverage of any of their contained summits 0x, 1x, or 2x the medianmax(medCov) across all peaks. The 0-median filtered set of peaks contained 859,563 peaks with a similar size distribution (notably, this step filtered some peaks as they had a median of 0 TMM across individuals at all of their summits, indicating most individuals did not have the peak); the 1-median filtered set of peaks contained 546,476 peaks with approximately two-fold enrichment in smaller peaks; and the 2-median filtered set contained 187,046 peaks with approximately 5-fold enrichment at smaller peaks (Supplemental Figure 7B&C). To determine if this filtering process removed expected true ATAC peaks, we examined the chromatin states each summit lied in, and performed a Fisher’s exact test on the remaining summits vs the filtered summits (Supplemental Figure 7D). We found that, across all filtering, the remaining peaks were enriched for promoters (up to 8-fold), enhancers (up to 3-fold), and bivalent chromatin (up to 22-fold), with larger enhancer enrichment in the 0- and 1-median filtered sets compared to the 2-median filtered set, and the inverse for promoters. Due to the large number of peaks removed by the 2-median filter, the enrichment of small peaks in the 1-median filter, and similar chromatin enrichment profiles (with 1-median filtering leaning toward enhancer enrichment), we chose to filter the peaks with the 1-median threshold, producing a final set of 546,476 peaks with a median size of 221bp (Supplemental Figure 7E), and 1,215,376 summits for analyses.

### H3K27AC ChIP-seq experiments and peak calling

We performed H3K27AC ChIP-seq in 52 iPSC samples from 46 lines from 36 individuals from iPSCORE. Pellets of formaldehyde -crosslinked iPSCs were lysed and sonicated in 110 µl of SDS Lysis Buffer (0.5% SDS, 50mM Tris-HCl pH 8.0, 20mM EDTA, 1x cOmplete™ Protease Inhibitor Cocktail (Sigma)) using Diagenode Bioruptor UCD-200 (Diagenode) or Covaris E220 Focused-ultrasonicators (Covaris). For each sample, 1 µg of H3K27ac antibody (Abcam ab4729) was coupled for 2-4h to 11µl of Protein G Dynabeads (Thermo Scientific) and used for overnight chromatin immunoprecipitation in IP buffer (1% Triton X-100, 0.1% DOC, 1x TE buffer, 1x cOmplete™ Protease Inhibitor Cocktail). 40-45µg of chromatin (66ug – 1 sample) was used for immunoprecipitation. Beads with immunoprecipitated chromatin were washed five times with 150 µl of RIPA buffer (50mM HEPES pH 8.0, 1% NP-40, 0.7% DOC, 500mM LiCl, 1mM EDTA, 1x cOmplete™ Protease Inhibitor Cocktail) and once with 1X TE buffer (10mM Tris-HCl pH 8.0, 1mM EDTA). Next samples were eluted in 150 µl of ChIP Elution Buffer (1% SDS, 10mM Tris-HCl pH 8.0, 1mM EDTA) and reverse crosslinked by incubation for overnight at 65°C and subsequent incubation with 5 µl RNAse (Sigma) for 1h at 37°C and Proteinase K Solution (20 mg/mL, Thermo Fisher Scientific) for 1h at 55°C. After reverse crosslinking, samples were purified with MIniElute PCR purification kit (Qiagen) or with DNA Clean & Concentrator kit (Zymo), eluted in 25µl of EB buffer (Qiagen)and Qubit (Thermo Scientific) quantified. Libraries were generated using KAPA Hyper Prep Kit (KAPA Biosystems) and KAPA Real Time Library Amplification Kit (KAPA Biosystems) at Institute for Genomic Medicine at University of California, San Diego. Libraries were barcoded using TruSeq RNA Indexes (Illumina). Libraries were sequenced on an Illumina HiSeq 4000 100bp Paired-End reads. Peaks were called using MACS2 --broadpeak on all samples simultaneously.

### ATAC-seq tag distribution at H3K27ac peaks

MakeTagDirectory.pl from HOMER v4.7^47^ was used on each sample individually at the set of H3K72ac peaks called on all samples simultaneously. After creating tag directories, annotatePeaks.pl from HOMER was used with -size 1000 -hist 50 -d to find mean coverages per sample at H3K27ac peaks. The average of this coverage was then calculated for plotting.

### Chromatin state enrichment at ATAC-seq peaks

To measure the enrichment for particular chromatin states at peaks, we used bedtools^48^ to identify the number of base pairs present in each chromatin state within ATAC peaks, and compared this proportion to that of the coverage of each state in the entire genome via a Fisher’s Exact test.

### Enriched transcription factors at accessible sites

To calculate the enrichment for transcription factors for sites, findMotifsGenome.pl from HOMER was used on hg19 with -size 200.

### Transcription factor motif prediction

To identify transcription factor (TF) binding sites at sites, FIMO from MEME v4.12.0^49^ was used on the 200bp flanking each sites with transcription factors from HOCOMOCO individually. Results were filtered to q<0.05 for each TF.

### Identifying co-accessible sites

To identify co-accessible sites, we utilized a Linear Mixed-effects Model (LMM) to control for fixed covariate effects, and random effects from kinship, as iPSCORE contains related individuals. First, we quantile normalized TMMs for each accessible site across individuals to remove outlier effects. Next, we utilized Limix v1.0.17 (github.com/limix/limix) and included age, iPSC passage number, sex, and the top 20 PCs from ancestry (previously calculated in Panopolous et. al^20^) as fixed effect covariates, and kinship as a random effect. Kinship values were obtained from DeBoever et. al^37^. We then ran Limix with these covariates on all pairs of sites for each chromosome, and Benjamini-Hochberg FDR corrected the regression p-values within each chromosome using statsmodels in python, using a significance threshold of q < 0.05. To examine within-peak correlation, we used the p-values from these analyses.

### Co-accessibility enrichment at co-binding TFs and chromatin looping

To measure co-accessible enrichments for TFs and chromatin looping, pgl files were created by pairing together EZH2 and Suz12 peaks within 500bp of one another, and pgltools^50^ was used to convert calls from CTCF ChIA-PET^34^ in GM12878 and pHiC in iPSCs^33^ to the pgl format. Next, pgltools intersect1D was used to find accessible sites at opposing anchors of loops, or at Ezh2 Suz12 pairs, and p-values were obtained from the co-accessibility analysis for enrichments.

### Annotating accessible sites with TF ChIP-seq, gene promoter, and chromatin states

To label TFs, gene promoters, and chromatin states at accessible sites, we utilized all public ChIP-seq from UCSC genome browser in ESC, ROADMAP chromatin state E020 15 state model, and GENCODE promoters. Bedtools was used to identify annotations which overlapped sites, and each site was labelled with all ChIP types, a chromatin state, and a gene (if it overlapped one).

### Annotating accessible sites with TF Motifs

Sites were annotated with all motifs they overlapped from the above FIMO analysis using bedtools. Following, sites were filtered for the clustering analysis by: 1) finding the mean TPM of all genes in the 134 individuals; 2) identifying enriched motifs with HOMER; 3) mapping HOCOMOCCO motif names to GENCODE genes using HOCOMOCCO’s metadata information (note: many genes were lost in this process); filtering to TFs whose genes were expressed at log_2_(TPM)≥1.

### Creation of co-accessibility networks

To create co-accessibility networks, we utilized the networkX package for python (v2.1). Edges were added to the network between each FDR q < 0.05 co-accessible sites with weights equal to their regression coefficient. All sites within the network were then annotated with chromatin states, gene promoters, TF ChIP-seq, and TF motifs, and edges were annotated with the genomic distance between sites. Following, all networks were combined into a single network for ease of use. Chromosome networks are induced by taking the subset of nodes within a given chromosome, and networks centered on a node are induced by subsetting to the node and all its neighbors.

### Clustering of annotations in co-accessibility networks using permutation tests

To create null networks, node labels (ie site names) were shuffled 25k times, and all annotations were shuffled with them; edges remained constant. To calculate empirical p-values, for each of the 25k null permutations, the mean edge weight between any two annotations was calculated and compared to the true mean edge weight. Directional empirical p-values were calculated by counting the number of times a stronger mean weight occurred in the null network compared to the original network, using the sign of the true mean edge weight (ie, as 1_Tss and 10_TssBiv had a positive mean weight, we counted the number of times a larger positive number occurred; as 3_TxFlnk and 15_Quies had a negative mean weight, we counted the number of times a larger negative number occurred). We then calculated an empirical p-value, and signed the p-value by the original mean edge weight sign so that anti-correlations would repel positive correlations for the same TF or chromatin state during clustering (ie TF_A and TF_B positive enrichment should cluster far away from an antagonistic association with TF_C).

### Identifying caQTLs and caSites

caQTLs were identified using the qtl_test_lmm function from Limix v1.0.17. TMM data was quantile normalized within each site across individuals. A kinship matrix was included as a random effect to account for relatedness between individuals, and the following variables were included as fixed effect covariates: age, sex, the top 20 principle components for ancestry, and the top 30 PEER factors from the TMM normalized count data (calculated with PEER v1.0^51^). While the ATAC samples were each sequenced twice, as each individual included data from both runs, batch was not included as a covariate. All sites were tested as we previously filtered our site set based on coverage (see above in methods). All SNVs within 100kb upstream or downstream of each site were utilized for testing. As the data space was large (∼1 million sites) we chose a conservative correction approach that was computationally fast: from the p-values for each SNV calculated from Limix, the minimum p-value was chosen from each site and Bonferroni corrected for the within-site number of tested SNVs. These Bonferroni adjusted p-values were then FDR-corrected as a whole across all sites, and sites with an FDR q-value < 0.05 were identified as significant caSites.

### Identifying allele specific effects at caSites

To identify allele specific effects (ASE), BAMS were remapped with WASP (v0.2.1), following which all heterozygous variants within 200bp upstream or downstream of a site were utilized. At each site, samtools^52^ mpileup was utilized to obtain allele counts. To identify ASE, all variants with 10 or more reads were tested for imbalance via a normal approximation to a binomial so that Z scores could subsequently be combined across individuals in a signed manner via Stouffer’s method. P values were calculated for each Stouffer Z score, and ASE sites were identified as those with one variant with an FDR q < 0.05.

### Concordance in ASE across co-accessible sites

To determine if ASE was similar across co-accessible sites, each node in the co-accessibility network was labelled with its ASE Z score. Next, we identified all pairs of Z scores connected by an edge. Finally, we utilized statsmodels.OLS.fromformula to regress the weaker Z scores against the stronger Z scores with a forced intercept of 0 (as no ASE would correspond to no ASE).

### Trans caQTL

To identify putative *trans-*caQTLs, we performed targeted association tests by leveraging the co-accessibility networks. For each *cis-*caSite, we tested the lead variant of the site against its co-accessible partners, and the FDR q-corrected all trans tests simultaneously with the Benjamini-Hochberg method. For these associations, we included sex, age, passage, the top 5 PCs from ancestry, and the top 20 PEER factors from expression as fixed effects, and kinship as a random effect.

### Mediator analysis for trans caQTL

To identify *trans* ca-QTLs whose effects were mediated through *cis-*caSites, we calculated the Sobbel p-value for each variant-cis-trans combination that was FDR q< 0.05 from the putative trans analysis. First, the trans height (Y_trans_) was regressed against the cis-site height (Y_cis_), using the SNP genotypes (X_cis_) as a covariate to obtain the mediator effect (β_mediated_) and its standard error 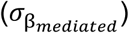:

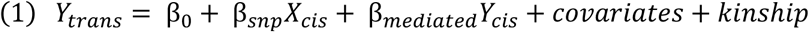

Next, the *cis* association (β_cis_) and its standard error 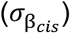 was obtained from prior analysis. The Sobbel p-value was found using the following Z score:

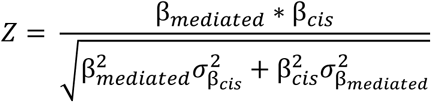

We utilized Limix to perform the analysis, and included the *cis* genetic effect as a fixed effect covariate in order to obtain the necessary values as output from Limix.

### Validation of GTEx trans-QTLs for co-accessibility

To identify if GTEx trans-QTLs had co-accessibility, we identified all sites at the 32 gene promoters, and all sites within 100kb of the reported eQTL variant (the converse of how we define what variants to test as caQTLs). We then tested all promoter sites against all variant sites using Limix and the methods described in the co-accessibility section. P-values were Bonferroni corrected within each eQTL-eGene pair, and then FDR corrected across all eGenes. A threshold of q < 0.05 was used for significance.

### Trans eQTL

To identify *trans* eQTLs, we tested *trans-*caQTL variants against the gene whose promoter overlapped the *trans-*caSite, following which we FDR corrected all eQTL tests. For these associations, we included sex, age, iPSC passage number, the top 5 PCs from ancestry, and the top 20 PEER factors from expression as fixed effects, and kinship as a random effect.

